# Genomic Engineering of Gene Dosage: A Generalizable Framework for Modeling Haploinsufficiency-Mediated Human Disorders through Splicing Modulation

**DOI:** 10.64898/2026.09.20.753023

**Authors:** Eric D. Smith, Joshua K. Meisner, Abbey Bullard, Connor B. Ward, Jennasea Licata, Cathy Smith, Colleen M. Kelly, Zachary T. LaRochelle, Sabrina Friedline, Aaron Renberg, Riley Yucius, Jesse Lange, Elizabeth D. Hughes, Thomas L. Saunders, Zachary T. Freeman, Yao-Chang Tsan, Michael J. Previs, Jacob Kitzman, Adam S. Helms

## Abstract

Heterozygous loss-of-function variants causing gene dosage reduction underlie many human genetic disorders, yet preclinical mouse models frequently fail to recapitulate human disease phenotypes due to post-translational compensation. Here, we present a generalizable framework to modulate gene dosage by alternative splicing via genome editing. By shifting proportions of transcripts toward nonsense-mediated decay, this approach enables precise titration of functional protein levels. Applying this concept to *Mybpc3*, we combined hypomorphic splice-altering alleles to overwhelm post-translational buffering in mice, successfully reproducing hallmark structural and functional features of human hypertrophic cardiomyopathy. Extending this concept, we engineered a portable Modulated Alternative Splice Cassette (MASC) inserted into *Dsp*, yielding patient-level protein reductions and characteristic arrhythmogenic cardiomyopathy pathologies, including subepicardial cardiac fibrosis and immune cell infiltration. These novel haploinsufficient models of *Mybpc3* hypertrophic cardiomyopathy and *Dsp* arrhythmogenic cardiomyopathy will critically enable mechanistic and therapeutic testing under protein stoichiometry conditions that closely replicate the patient disorders. Finally, systematic *in silico* predictive modeling across 645 human haploinsufficiency-associated genes demonstrated the generalizability of MASC insertion effects on splicing across diverse tissue types. Notably, the framework developed here leaves endogenous gene expression regulatory logic intact, allowing these models to be used to develop and validate therapeutic approaches that target transcription. Overall, this strategy provides a scalable roadmap for engineering high-fidelity animal models for human diseases caused by haploinsufficiency.

## Introduction

Most novel medications fail during clinical development, often due to lack of efficacy in human clinical trials.^1–3^ In most cases, medications with lack of efficacy in humans have been previously tested in preclinical animal model studies that had shown apparent benefit. This discrepancy is presumably due to biologic differences in animal models, including how well the models resemble the human disease phenotypes. In a systematic study from the International Mouse Phenotyping Consortium (IMPC), only ∼50% of mouse gene mutation models recapitulated the human ortholog disease phenotypes.^4^ A large contributor to this discordance was found to be differential tolerability of heterozygous states in mice compared to humans.^4^ Generally, mice have been shown to be more resilient to gene dose effects, hindering the development of accurate models of human diseases.^4^

Single allele loss of function variants that cause reduced protein levels (i.e., haploinsufficiency) are collectively a common cause of disease in humans. Current curated lists from the Clinical Genome Resource (ClinGen) include 460 genes with sufficient evidence for haploinsufficiency.^5^ Quantitative studies of single allelic loss of function tolerance in large human populations revealed an even larger burden of haploinsufficiency genes, approximately 3000, with heterozygous loss of function variants in these genes accounting for a broad variety of human disorders.^6^ Thus, generation of accurate murine models of these disorders, recapitulating the dosage sensitivity observed in humans, is critically important.

Heart muscle disorders (cardiomyopathies) are most frequently caused by autosomal dominant variants in genes affecting myofilament contraction and/or structural coupling in cardiomyocytes. Haploinsufficiency has been shown to be the most common mechanism among prevalent genetic subtypes of cardiomyopathy. For example, the most commonly associated gene for hypertrophic cardiomyopathy (HCM) is *MYBPC3*, encoding cardiac myosin-binding protein C (cMyBP-C), which regulates actin-myosin interactions in the cardiac sarcomere.^7–11^ Most patient variants in *MYBPC3* result in loss of function and haploinsufficiency.^8,12–14^ The most common gene associated with the left ventricular dominant form of arrhythmogenic cardiomyopathy (ACM) is *DSP*, encoding desmoplakin, which links desmosomes to the intermediate filaments.^15^ For these and other key cardiomyopathy-associated genes (e.g., *TTN*, *LMNA*, *PKP2*), mouse models with heterozygous loss of function variants that cause cardiomyopathy in patients fail to recapitulate robust cardiomyopathy phenotypes under baseline physiologic conditions.^16–22^ This discordance is likely driven in part by post-translational dosage compensation, wherein cellular degradation pathways adjust turnover rates to maintain protein stoichiometry despite reductions in transcript abundance.^23,24^ As a result, investigators have resorted to homozygous knock-out models.^17,22,25,26^ While informative for probing gene function, these models have limited preclinical fidelity, since complete loss of function phenotypes are not simply exaggerated versions of partial dosage reduction – the underlying phenotypes may diverge fundamentally.

Here, we developed an approach for generating haploinsufficient models with graded reductions in gene dose with the goal of replicating human levels of haploinsufficiency and disease phenotypes in mice. To leave endogenous transcriptional promoter and enhancer regulation intact, we reasoned that alternative splicing could be leveraged to shift a proportion of transcripts toward nonsense mediated decay (NMD) sensitive transcripts resulting in partial null (hypomorphic) alleles. We first applied this approach to *MYBPC3*, for which we identified a hypomorphic splice gain variant in an HCM patient population.^27^ Using SpliceAI to predict alternative splicing effects in the murine context, we generated a panel of alternative splice-susceptible alleles in mice.^28^ We measured both mRNA and protein levels, demonstrated their nonlinear relationship, and validated the allelic expression reduction needed to generate patient-levels of haploinsufficiency. The resultant haploinsufficient *Mybpc3* mouse model exhibited key features of human HCM. We then extended this approach to design a Modulated Alternative Splice Cassette (MASC) that was inserted upstream of a constitutively expressed exon in *Dsp*. Multiple versions of the splice cassette, predicted to have graded alternative splicing by SpliceAI, were introduced concurrently using pooled homologous recombination knock-in. Similarly, mRNA and protein levels were quantified for multiple allelic combinations to validate the mRNA dose reduction needed to generate haploinsufficiency. The resultant model exhibited patient levels of haploinsufficiency and key features of human ACM. Finally, we performed systematic *in silico* estimations of graded effects with alternative splice cassette insertion for a comprehensive list of haploinsufficiency-associated genes, suggesting portability of this concept across tissue types and providing a roadmap for generation of haploinsufficient genetic models for a broad array of human disorders.

## Results

### A hypomorphic splice variant in *Mybpc3* results in an additive gene dosage effect to recapitulate haploinsufficiency

Null alleles, which are typically used to generate disease models, result in increments of either 50% (heterozygous) or 100% (homozygous) knock-out of mRNA transcripts. Compensatory mechanisms in mouse models are known to mitigate the effects of haploinsufficiency, as has been shown previously in the case of *Mybpc3* truncating variants.^16,19–22^ We verified this limitation for a previously published heterozygous *Mybpc3* truncating variant mouse model using a highly precise mass spectrometry assay of cMyBP-C protein levels, benchmarked against prior data from the same assay in human patient samples (**Figure 1A**).^14,29^ In contrast to heterozygous truncating variants in patients, the heterozygous mouse model exhibited no significant decrement in cMyBP-C level, while homozygous truncating variants resulted in a complete loss of cMyBP-C.

**Figure 1.**
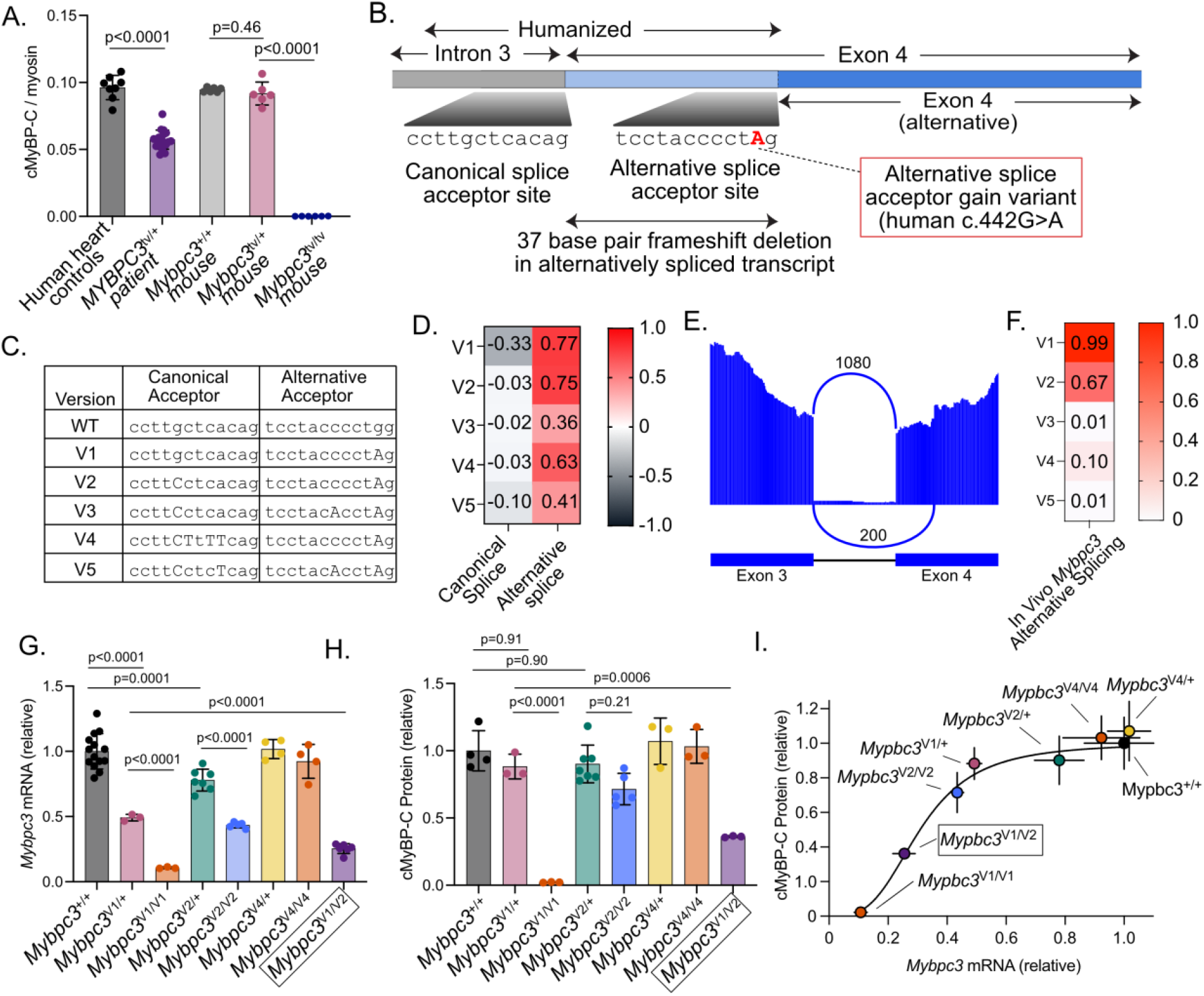
A hypomorphic splice variant in *Mybpc3* results in an additive gene dosage effect to recapitulate haploinsufficiency in mice. **A.** Mass spectrometry was performed to precisely compare cMyBP-C levels of heterozygous and homozygous *Mybpc3* truncating variant mice compared to wild-type mice. No reduction in cMyBP-C level was observed in *Mybpc3*^tv/+^ mice, in contrast to measurements of heart samples from *MYBPC3*-HCM patients with heterozygous truncating variants (as previously reported in O’Leary, et al.). **B.** The *MYBPC3* c.442G>A variant in humans results in a partial splice gain acceptor site that competes with the canonical splice site. This variant was introduced into mice along with the preceding portion of exon 4 and the intronic splice acceptor site. Transcripts utilizing the splice gain site have an effective 37 bp frameshift deletion, rendering these transcripts susceptible to NMD. **C.** Tested splice variant sequences are shown (variants capitalized). **D.** SpliceAI predictions are shown for each variant combination. Predicted splice acceptor site losses are shown as negative values (range: -1 to 0 for splice acceptor loss, 0-1 for splice acceptor gain). **E.** A representative Sashimi plot is shown, demonstrating partial alternative splicing at the exon 3-4 junction. **F.** Alternative splicing quantification for each variant measured from RNA-seq from mouse hearts. **G.** mRNA levels for each wildtype, heterozygous, and biallelic model are shown (mRNA levels quantified by RNA-seq). **H.** cMyBP-C levels for each wildtype, heterozygous, and biallelic model are shown (cMyBP-C levels quantified by mass spectrometry). I. *Mybpc3* mRNA and cMyBP-C protein levels were correlated for each model.

To overcome this limitation, we reasoned that a hypomorphic (partial null) allele could act as sensitizing alleles by causing only a partial reduction of transcript levels. Such an allele could then be combined to allow titration of the composite gene dose by overwhelming post-translational compensation. We recently identified and validated a hypomorphic splice variant in *MYBPC3* in patients with HCM – therefore, we selected this variant (*MYBPC3* c.442G>A) to initially test our hypothesis.^27^ In humans, *MYBPC3* c.442G>A introduces a missense variant (p.Gly148Arg) but the pathogenic effect of the variant has been demonstrated to be gain of a novel splice acceptor site that competes with the canonical splice acceptor site of exon 4.^27^ This alternative splice gain causes a frameshift and a corresponding total allelic transcript reduction of 41±13% due to NMD of the alternatively spliced transcripts.^27^

Since the mouse genomic sequence differs in exon 4, we performed genome editing of this site to simultaneously “humanize” this sequence while also introducing multiple intronic or synonymous variants to modulate splice competition between the canonical and splice acceptor gain site (**Figure 1B-C**). We chose the variants based on SpliceAI predictions of alternative splice site utilization to create multiple unique alleles (variant combinations V1-V5) that would allow total gene dose titration (**Figure 1C-D**). To efficiently generate multiple alleles in mice, we then performed pooled genome editing in mice by using an equimolar concentration of each repair template during CRISPR-Cas9 editing. All 5 intended edits were recovered among 125 G0 progeny.

We then analyzed alternative splicing for each allele by RNA-seq. Analysis of exon junctions demonstrated alternative splicing between the canonical and splice acceptor gain site, similar to the alternative splicing observed in human cardiomyocytes differentiated from induced pluripotent stem cells (iPSC-CMs, **Figure 1E**).^27^ As intended, a 37 base pair deletion was effectively introduced in alternatively spliced transcripts, shifting the reading frame and causing a downstream premature termination codon, rendering these transcripts susceptible to NMD. Three of the five variant combinations resulted in substantial proportions of alternative splicing – 99%, 67%, and 10% for V1, V2, and V4, respectively (**Figure 1F**). No detectable alternative splicing was present for the V3 and V5 alleles.

Next, we generated heterozygous and biallelic mouse models that we predicted would cause a broad range of mRNA level reduction by using combinations of the V1, V2, and V4 splice gain alleles. *Mybpc3* mRNA and cMyBP-C protein levels were measured by RNA-seq and mass spectrometry, respectively. Across this series of models, *Mybpc3* mRNA and cMyBP-C protein levels varied in a graded manner, with V4/+, V2/V2, and V4/V4 genotypes producing intermediate reductions in mRNA that did not significantly lower cMyBP-C protein levels (**Figure 1G-H**), consistent with continued post-translational buffering at these more modest dosage reductions. The allele with the highest proportion of alternative splicing (i.e., V1, 99%) in heterozygous state caused a 51±3% reduction in *Mybpc3* mRNA, consistent with this allele being effectively null. As with prior heterozygous null alleles (**Figure 1A**), this allele in heterozygous state did not alter total cMyBP-C level (**Figure 1H**). However, combining this allele with the V2 allele (i.e., the *Mybpc3*^V^^1/V^^2^ biallelic model) resulted in a 64±1% reduction in cMyBP-C (**Figure 1H**), more closely matching cMyBP-C reductions in heterozygous HCM patients (**Figure 1A**). Plotting total mRNA transcript levels vs. cMyBP-C protein levels for all of the tested models revealed a non-linear, sigmoidal dose-response relationship (fit to a model constrained to biological boundaries of zero and 1: Hill slope=3.238, EC50=0.309, R²=0.88). This relationship indicated that cMyBP-C protein levels are relatively preserved across a broad range of mild-moderate mRNA reduction, with more substantial protein loss occurring only once mRNA levels fall below approximately 30% of normal. Taken together, these data demonstrate that post-translational compensatory mechanisms that maintain protein level stoichiometry can be overwhelmed by additive effects from hypomorphic splice variants to attain haploinsufficiency in mice.

### Haploinsufficiency of cMyBP-C causes hypertrophic cardiomyopathy in mice

We next characterized the phenotype of the haploinsufficient *Mybpc3*^V^^1/V^^2^ mice. These mice were generated by breeding *Mybpc3*^V^^1/+^ with *Mybpc3*^V^^2/V^^2^ mice to generate a 50:50 ratio with *Mybpc3*^V^^2/+^ littermate controls, the latter having normal levels of cMyBP-C, as shown above (**Figure 1H**). Left ventricular mass calculated from echocardiography was increased by 1 month of age (**Figure 2B**). Left ventricular ejection fraction (LVEF) was not significantly different (**Figure 2C**). Isovolumic relaxation time, a sensitive measure of diastolic function in mice, was significantly prolonged, consistent with human *MYBPC3* HCM (**Figure 2D**).^30,31^ Aortic ejection time, a parameter capable of detecting subclinical systolic impairment before changes in ejection fraction, was reduced in haploinsufficient mice (**Figure 2E**), also consistent with human *MYBPC3* HCM.^32^ Additionally, mouse hearts were weighed directly at time of dissections – heart weight to body weight ratios were significantly increased in haploinsufficient mice by 1 month of age, persisting through 4 months of age (**Supplemental Figure 1A**). We also characterized *Mybpc3*^V^^4/V^^4^ mice, which have no significant decrease in cMyBP-C protein but exclusively express the humanized portion of exon 4 including the c.442G>A variant. These mice did not exhibit hypertrophy or echocardiographic evidence of functional deficits, excluding the possibility that either the humanized exon 4 sequence or c.442G>A variant, independent of splicing effects, is sufficient to have contributed to the phenotype observed in the haploinsufficient model (**Supplemental Figure 1B-D**).

**Figure 2.**
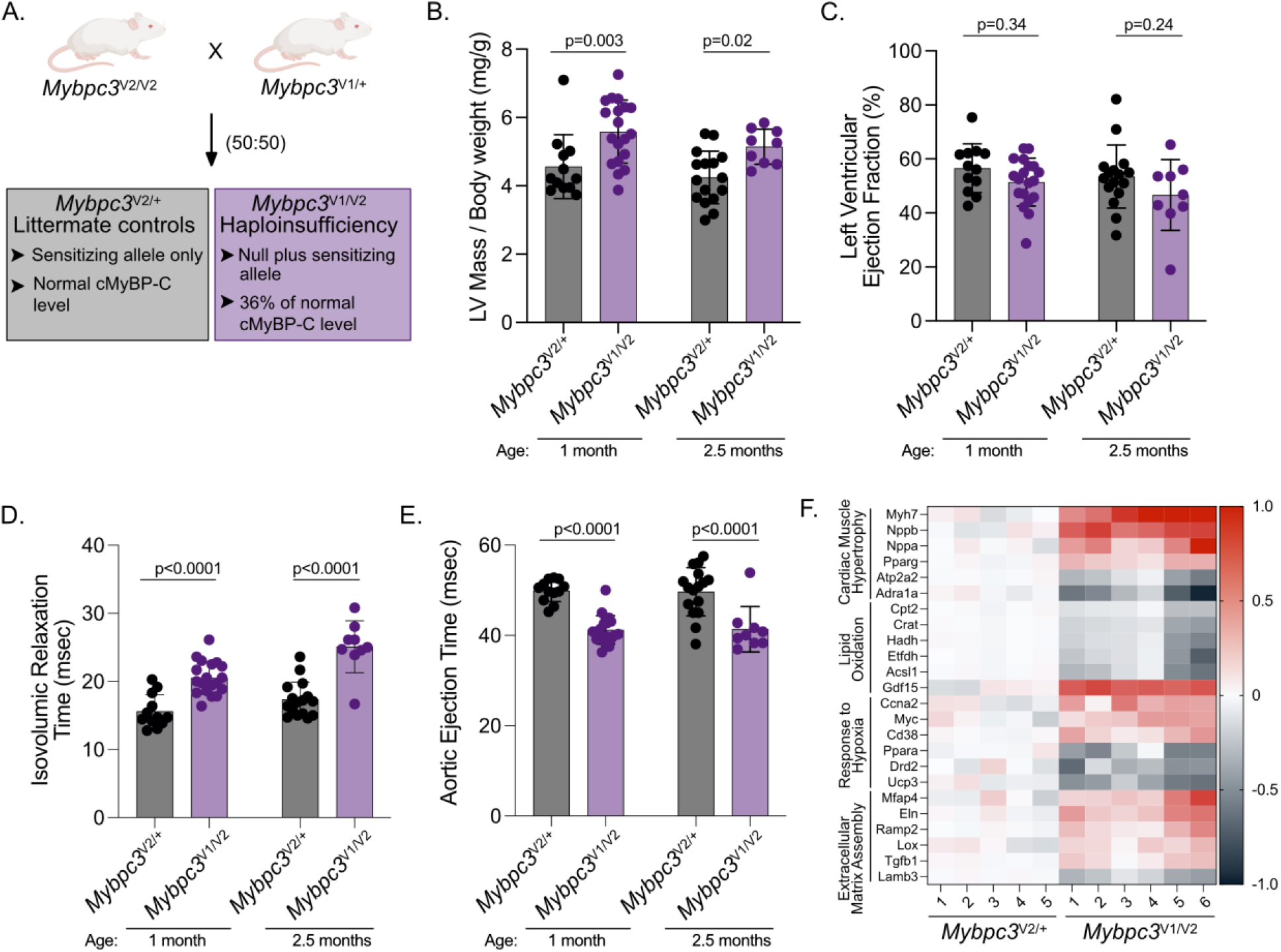
Haploinsufficient *Mybpc3* mice develop cardiac hypertrophy and functional deficits. **A.** To generate biallelic mice, *Mybpc3*^v^^2/v^^2^ mice (homozygous for the hypomorphic V2 allele) are bred with *Mybpc3*^v^^1/+^ mice (heterozygous for the ∼null V1 allele) to generate litters with 50:50 biallelic, haploinsufficient *Mybpc3*^v^^1/v^^2^ mice. *Mybpc3*^V^^2/+^ mice have normal cMyBP-C protein levels and serve as littermate controls. **B-E.** LV mass, LV ejection fraction, aortic ejection time, and isovolumic relaxation time were assessed by echocardiography and compared at both 1 and 2.5 months of age. **F.** Transcriptional analyses were performed for *Mybpc3*^v^^1/v^^2^ (N=6) vs. *Mybpc3*^V^^2/+^ (N=5) mice at 4 months of age by RNA-seq. Differentially expressed genes for representative dysregulated biological pathways are shown with each column representing an individual biological replicate.

We assessed transcriptional patterns of cardiac remodeling by RNA-seq at 4 months of age. Differentially regulated genes are shown in **Supplemental Figure 2A**. We performed gene set enrichment (GSEA) and gene ontology analyses, revealing dysregulated biological pathways characteristic of human HCM, including “cardiac muscle hypertrophy in response to stress” (p=0.002), “lipid oxidation” (p=9.2e-4), “response to hypoxia” (p=0.002), and “extracellular matrix assembly” (p=8.1 e-4).^29,33^ Relative expression levels for representative genes for these pathways are shown in **Figure 2F** and top dysregulated gene sets from GSEA are shown in **Supplemental Figure 2B**. Expression levels of *Nppb* and *Myh7*, key markers of cardiac stress and adverse remodeling, were markedly increased in *Mybpc3*^V^^1/V^^2^ mice (**Supplemental Figure 2C-D**). These markers were not different in *Mybpc3*^V^^4/V^^4^ mice (**Supplemental Figure 2E-F**).

### Generation of a modulated alternative splice cassette (MASC) allows control of *Dsp* expression level in vivo

Similar to *MYBPC3*, heterozygous truncating variants in *DSP* are a major cause of cardiomyopathy in patients through a loss of function mechanism.^15,34^ We first obtained a global *Dsp* knock-out model from the Knock-Out Mouse Project (KOMP) repository to determine the heterozygous truncating variant phenotype.^35^ The KOMP *Dsp* model contains an exon 3 deletion that results in a frameshift with consequent premature termination codon. We found that *Dsp^Ex^*^3d^*^el/+^* mouse hearts exhibited reduced mRNA expression levels to 42 ± 4% of wild-type (p<0.0001; **Figure 3A**). However, Dsp protein level was reduced more modestly (68 ± 10% of wild-type), consistent with partial compensation (p=0.02, **Figure 3B-C**). Additionally, *Dsp^Ex^*^3d^*^el/+^* heterozygous mice did not develop excess myocardial fibrosis, the defining feature of ACM (histological quantification of extracellular matrix: 7.4 ± 1.4% vs 7.2 ± 3.4%, p=0.86, **Figures 3D-E**). These results demonstrated partial gene dosage compensation and lack of a clear disease phenotype in heterozygous *Dsp* knock-out mice.

**Figure 3.**
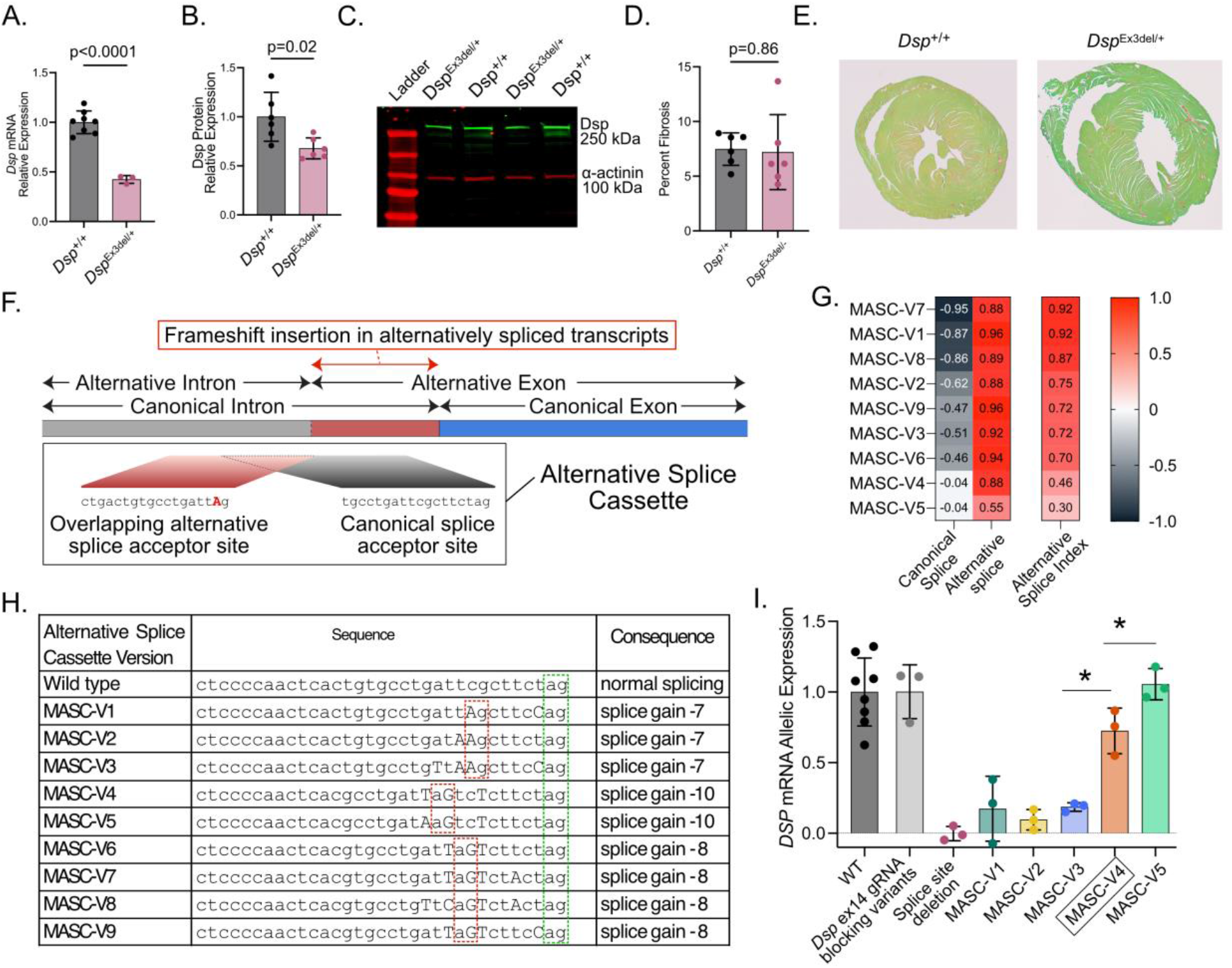
Generation of a modulated alternative splice cassette (MASC) allows titration of Dsp expression in vivo. **A-C.** Heterozygous *Dsp* truncating variant mice (*Dsp*^Ex3d^^el/+^) express 42 ± 4% of wild-type mRNA level (A) 68 ± 10% of normal protein level (B-C), consistent with partial post-translational compensation. **D-E.** *Dsp*^Ex3d^^el/+^ mice do not exhibit significant cardiac fibrosis, as quantified by picrosirius red staining. **F.** The modulated alternative splice cassette (MASC) was designed to create graded splice competition between the canonical splice acceptor site and an upstream out-of-frame splice acceptor site. **G.** Potential splice competition cassettes were evaluated by SpliceAI to identify variant combinations with a range of predicted effects on alternative splice site usage. Predicted effects of each MASC variant sequence inserted upstream of *Dsp* exon 14 on canonical splice acceptor loss and alternative splice acceptor gain are shown. The alternative splice index was defined as the average of the absolute values for splice acceptor gain and loss. **H.** The specific MASC sequence variants and positions of the alternative splice gain 5’ AG sites (dotted red boxes) and the canonical splice 5’ AG sites (dotted green box) are shown. Capital letters indicate variants compared to the wild type *Dsp* exon 14 splice acceptor sequence. **I.** Allelic mRNA expression levels for each MASC variant combination were quantified by RNA-seq. Expression levels are also shown for mice with a deleted splice site and with gRNA protospacer blocking variants that were positioned upstream of the splice acceptor sequence.

Therefore, with a goal of generating a *Dsp* model that more closely approximates the level of DSP protein haploinsufficiency observed in patients^34^, we extended the concept of alternative splicing modulation that we had implemented for *MYBPC3*. Additionally, we aimed to develop and validate a generalizable approach to titrate gene dosage through splice modulation that would be broadly applicable to haploinsufficiency-driven human disorders. To accomplish this goal, we hypothesized that SpliceAI based predictions could guide the introduction of hypomorphic alternative splice acceptor gain sites upstream of canonical splice acceptor sites, allowing titratable splice site usage without affecting coding regions. We selected the *Dsp* exon 14 splice acceptor site as a starting point since exon 14 is constitutively spliced in *Dsp* and follows a typical splice acceptor consensus sequence (i.e., U2-type with polypyrimidine tract followed by AG at the -2 to -1 positions). Using SpliceAI, we tested combinations of variants at this location that would create putative splice acceptor sites at upstream locations with consequent disruption of *Dsp*’s reading frame (**Figure 3F**). We refer to the 33 bp sequence comprising the two competing splice sites as the “modulated alternative splice cassette” (MASC). We identified nine different variant combinations that were predicted to use each of the alternative and canonical splice acceptor sites within the cassette to variable extents (**Figure 3G-H**). We defined an “alternative splice index” as the average of the absolute values for the SpliceAI predicted canonical splice acceptor loss and alternative splice acceptor gain (**Figure 3H**).

We then performed pooled homology-directed repair with CRISPR-Cas9 editing to introduce alternative splice cassette variants into mice. We selected 5 of the MASC variants for in vivo testing. Following CRISPR-Cas9 editing with equimolar concentrations of these repair templates, we recovered all 5 of the MASC variants, in addition to alleles containing gRNA blocking edits only and a canonical splice site deletion, among 96 G0 progeny. RNA-seq analysis revealed that MASC variants resulted in a range of effects on alternative splice site usage and consequent relative allele expression (**Figure 3I**). The effects of each MASC variant on allelic expression correlated with the SpliceAI alternative splice index (R^2^=0.88, p=0.01 vs non-zero slope).

### Biallelic splice modulation results in Dsp haploinsufficiency in mice

To determine *Dsp* mRNA and protein correlations in mice, we crossed mice to generate single and combinations of variants expected to create a range of effects on Dsp protein level. Since *DSP* is expressed highly in the skin as well, we additionally generated a knock-out allele for only the cardiac-predominant transcript variant 1 (TV1), which accounts for 77% of *DSP* expression in the human heart.^34^ *DSP* TV1 truncating variants cause cardiomyopathy in patients indistinguishable from global *DSP* truncating variants.^15,36^ We reasoned that mouse models carrying the *Dsp*-TV1 knock-out allele would have reduced susceptibility toward skin defects that may confound future analyses. We recently created a similar model with iPSC-CMs by introducing a truncating variant in the TV1-specific portion of exon 23, and we followed the same strategy here.^34^ Although patients with homozygous *DSP* TV1 truncating variants have been reported with severe phenotypes^37,38^, we recovered no homozygous *Dsp*^TV1k^^o/TV1k^° mice among ≥50 pups from ≥5 litters, consistent with non-viability. Additionally, crosses among *Dsp*^TV1k^^o/+^, *Dsp*^Ex3d^^el/+^, and mice carrying MASC variants with strong splice disruption (MASC-V1, MASC-V2, and MASC-V3) yielded no biallelic progeny in at least 50 pups from at least 5 litters for each combination, consistent with non-viability. Extrapolating from allelic expression analyses (**Figure 3A**, **Figure 3I**), these results indicate that *Dsp* mRNA levels below ∼20% of normal are not tolerated during fetal development in mice. In contrast, mice carrying biallelic MASC-V4 and TV1 knock-out alleles (*Dsp*^V^^4/TV^^1k^°) were born in the expected Mendelian ratio (∼50:50) and exhibited no apparent skin or hair abnormalities. Across this series of models (Dsp^⁺/⁺^, Dsp^V^^4/+^, Dsp^TV1k^^o/+^, Dsp^Ex3d^^el/+^, and Dsp^V^^4/TV^^1k^°), *Dsp* mRNA and protein levels declined progressively, with Dsp^V^^4/TV^^1k^° mice exhibiting *Dsp* mRNA levels 39±6% of normal and Dsp protein levels 30±4% of normal (**Figure 4A-B**). As with *Mybpc3* (**Figure 1I**), plotting Dsp protein level as a function of mRNA level across this allelic series revealed a non-linear, sigmoidal dose-response relationship (**Figure 4C**, R²=0.997). However, the Dsp curve exhibited a steeper Hill slope (5.16 vs. 3.238 for *Mybpc3*) and a higher EC50 (0.46 vs. 0.31 for *Mybpc3*), indicating that Dsp protein levels are preserved across a narrower range of mRNA reduction before declining sharply, consistent with a comparatively more limited capacity for post-translational buffering. We recently reported a similar level of DSP protein reduction in DSP cardiomyopathy patients undergoing heart transplant using a similar mass spectrometry technique (48±23% of normal, range 23-68%).^34^

**Figure 4.**
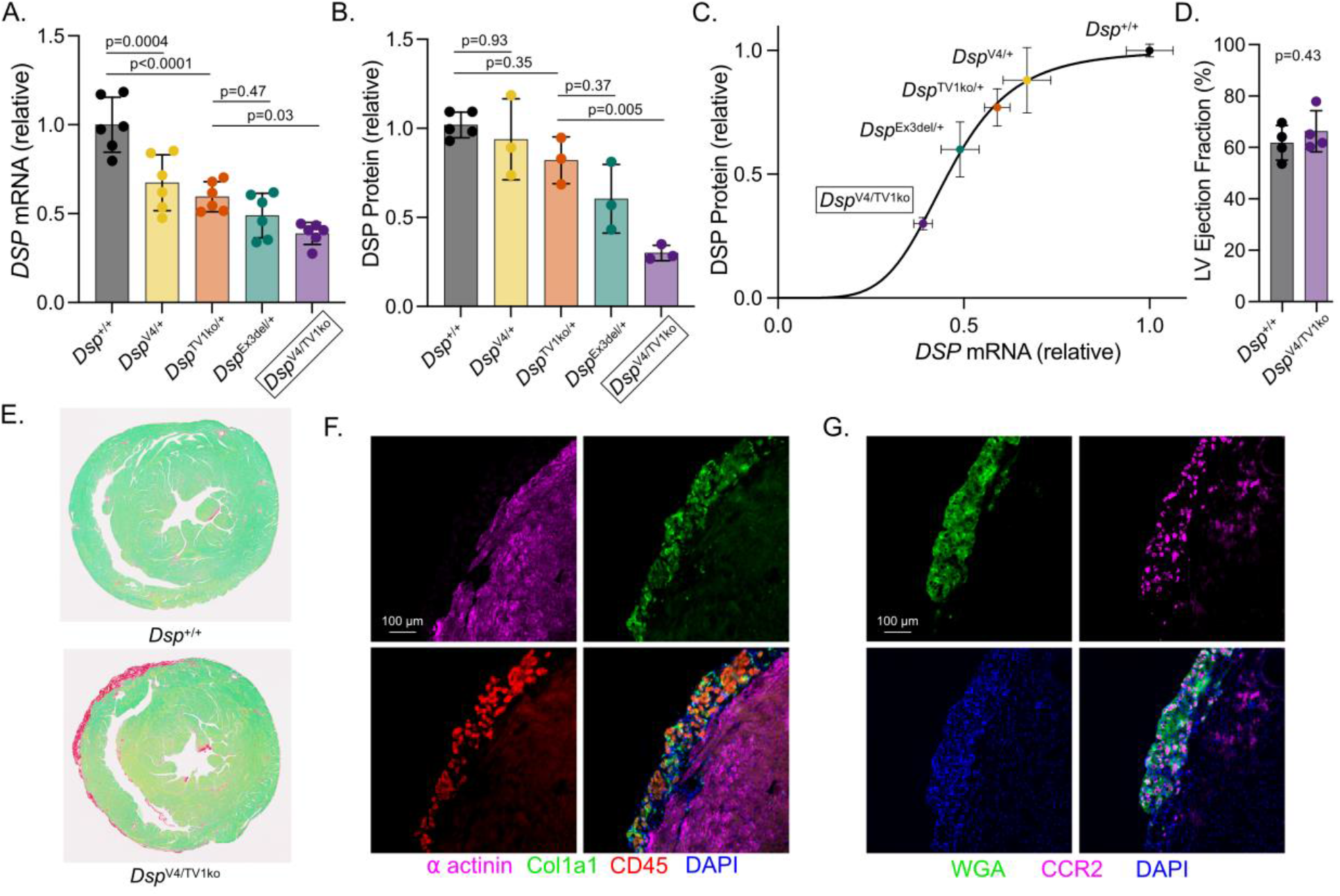
Biallelic splice modulation results in Dsp haploinsufficiency and cardiac fibrosis in mice. **A.** mRNA levels for wildtype, heterozygous, and biallelic models are shown (mRNA levels quantified by RNA-seq). **B.** Dsp protein levels for wildtype, heterozygous, and biallelic models are shown (Dsp protein level quantified by mass spectrometry). **C.** *Dsp* mRNA and protein levels were correlated for each model. **D.** LV ejection fraction was quantified by echocardiography at 3 months of age. **E.** Picrosirius red staining of cardiac cross-sections showed subepicardial fibrosis in *Dsp*^V^^4/TV^^1k^° mice but not in controls. **F.** Immunostaining of cardiac cross-sections was performed with antibodies for α-actinin, Col1a1, CD45, and DAPI. A representative section from a *Dsp*^V^^4/TV^^1k^° mouse is shown depicting a region of subepicardial fibrosis. **G.** Immunostaining of cardiac cross-sections was performed with antibodies for wheat germ agluttinin (WGA), CCR2, and DAPI. A representative section from a *Dsp*^V^^4/TV^^1k^° mouse is shown depicting a region of subepicardial fibrosis.

We next assessed the cardiac phenotype of haploinsufficient *Dsp*^V^^4/TV^^1k^° mice. The primary defining feature of DSP cardiomyopathy in humans is cardiac fibrosis, predominantly subepicardial fibrosis, which develops prior to overt systolic dysfunction.^15,39^ LV ejection fraction was preserved in 3-month-old *Dsp*^V^^4/TV^^1k^° mice (**Figure 4D**). Picrosirius red staining of 3-month-old *Dsp*^V^^4/TV^^1k^° whole heart sections revealed fibrotic remodeling specifically in the subepicardial muscle layer that is absent in control hearts (**Figure 4E**). Subepicardial fibrosis was present in 9 of 9 *Dsp*^V^^4/TV^^1k^° mice vs 0 of 6 control mice (p=0.0002, Fisher’s exact). Immunostaining for Col1a1 and CD45 revealed co-localization of fibrosis and immune cells (**Figure 4F**). Additionally, staining for WGA and CCR2 also demonstrated co-localization, indicating the presence of CCR2+ macrophages as a potential exogenous driver of inflammation and fibrosis in response to cardiac injury (**Figure 4G**). These findings replicate the key pathologies of fibrosis and inflammation preceding systolic dysfunction that are characteristic of DSP haploinsufficiency-associated cardiomyopathy in patients.^15,34,39–42^

### Predictions of modulated alternative splice cassette effects in other haploinsufficiency-associated genes are stable across tissues

Having demonstrated the potential of the alternative splice cassette approach to modulate gene dose, we next assessed the potential of this approach for broad application across haploinsufficiency-associated genes. We first assessed potential utility for cardiomyopathy associated with the gene *TTN* (encoding titin). *TTN* truncating variants are the most common genetic cause of dilated cardiomyopathy. Mice with *Ttn* truncating variants, however, exhibit a negligible phenotype in the absence of additional stressors.^18^ We derived SpliceAI predictions for the insertion of MASC sequences replacing 33 bp 5’ of *Ttn* splice acceptor sites. We selected 8 different constitutively spliced exons (i.e., high percent spliced in, PSI) for which truncating variants have been associated with dilated cardiomyopathy in patients.^43,44^ In each of these locations, predictions of splice competition behavior were similar to the above predictions for *Dsp*, and MASC-V1 through V9 variants demonstrated graded predicted activity (**Figure 5A**).

**Figure 5.**
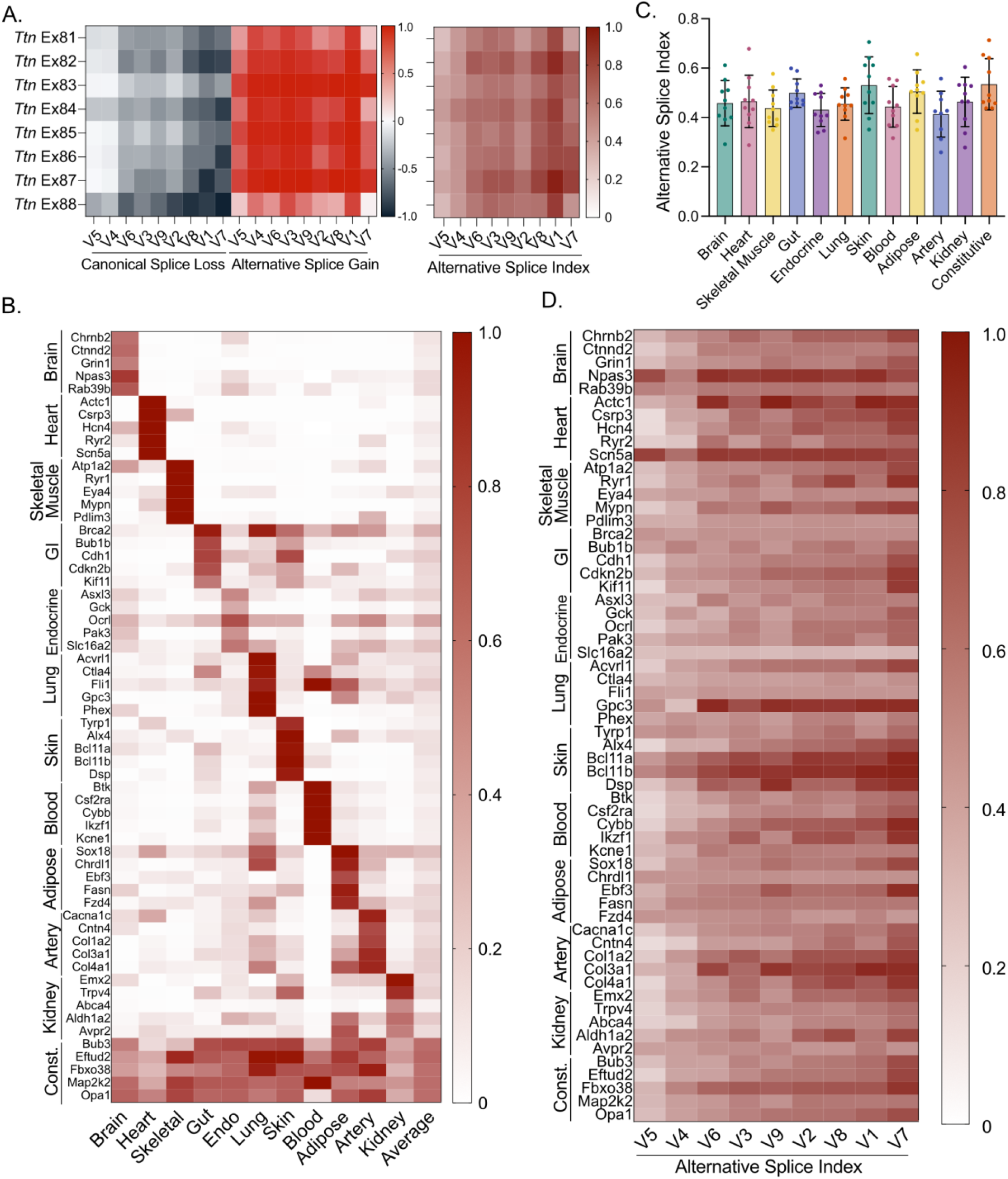
Predictions of modulated alternative splice cassette effects in other haploinsufficiency-associated genes are stable across tissues. **A.** SpliceAI predictions for canonical splice acceptor loss and alternative splice acceptor gain are shown for insertion of MASC-V1 through V9 variants, replacing the 33 bp sequence 5’ of the splice acceptor site for each of 8 exons of *Ttn*. The alternative splice index (as defined above) for these insertions is also shown (right). **B.** Genes with tissue specific expression profiles were identified from GTEx to allow comparison of MASC insertion effects. The expression profiles for the top 5 ranked tissue-specific genes for each tissue are shown, with expression scaled 0-1 based on the highest expressing tissue. **C.** Tissue-specific genes were compared for MASC-V4 insertion effects (ANOVA with Kruskal-Wallis tests for multiple comparisons were all non-significant). **D.** SpliceAI-derived splice index values demonstrated a range of predicted splice competition across tissue-specific and constitutively (“const.”) expressed genes.

We then calculated SpliceAI estimates of MASC effects for a comprehensive set of genes associated with haploinsufficiency-mediated disease in humans.^5,45^ Of this list of genes (N=707), we identified those with 1) mouse orthologs, 2) at least 3 exons, and 3) splicing data available in VastDB (N=645 human genes corresponding to 654 orthologous mouse genes).^46^ For each gene, two constitutively spliced exons (other than exon 1 or the terminal exon) with high PSI were selected for *in silico* MASC insertions that replaced the corresponding 33 bp sequence 5’ of the splice acceptor site (*N.B.*, only one exon was selected for genes with only 3 exons). SpliceAI predictions for all MASC variants for selected exons for each of the 654 genes are shown in **Supplemental Table 1**. We then assessed whether the SpliceAI MASC predictions were stable across genes expressed in different tissue types. Gene expression data from the Genotype-Tissue Expression (GTEx) database were extracted for the above set of haploinsufficiency-associated genes, which were then ranked for tissue-specific expression patterns (**Figure 5B**). Among the top 10 tissue-specific genes for each tissue type, the effects of MASC introduction were similar on average (as shown, for example, for MASC-V4 in **Figure 5C**). MASC insertion effects on alternative splicing exhibited some context dependent variability (**Figure 5D**). However, 118 of 120 (98%) of the tissue specific genes were predicted to have at least one MASC variant insertion effect in the splice index range of 0.4-0.7 for the selected exon. This range was most likely to result in an intermediate range of alternative splicing based on our in vivo results for *Mybpc3* and *Dsp*. Taken together, these analyses indicate that alternative splice cassette insertion has the potential to modulate gene dose across a broad range of genes associated with haploinsufficiency disorders.

## Discussion

A lack of accurate models of human haploinsufficiency disorders has been an obstacle to both mechanistic study and therapeutic development. Here, we have developed an approach based on hypomorphic splicing variants known to exert additive phenotypic effects in human genetic disorders. By modulating splicing, this approach creates stable gene expression levels that are capable of altering protein abundance along a continuum, rather than being limited to the all-or-none expression effects of knock-out alleles. Additionally, this approach avoids the heterogeneity, complexity, and limited titratability of other strategies, such as RNA silencing, CRISPR inhibition, or degradation sequences.^47–49^ Moreover, the MASC platform leaves native promoter/enhancer regulatory elements fully intact, making it uniquely suited for long-term physiological modeling, therapeutic testing, and evaluating locus-specific transcriptional targeting.

We initially developed our splice modulation approach for *Mybpc3*. Despite *MYBPC3* loss of function variants being the most common cause of hypertrophic cardiomyopathy, preclinical studies often avoid use of *MYBPC3* models because heterozygous truncating variants do not cause a clear disease phenotype. In contrast, homozygous truncating variants lead to a drastic fetal onset cardiomyopathy with unique mechanisms that lead invariably to infant mortality in humans.^25,50,51^ Our initial studies with the haploinsufficiency model revealed hypertrophy and diastolic dysfunction, characteristic of *MYBPC3*-HCM.^10,31^ Additionally, we identified evidence of systolic impairment through a reduction in aortic ejection time – this finding is notable in that patients with *MYBPC3*-HCM also exhibit a reduction in aortic ejection time, a distinct finding from *MYH7*-HCM.^32^ Additionally, the haploinsufficient *Mybpc3* model exhibited transcriptional evidence of metabolic and extracellular matrix remodeling, recapitulating key human *MYBPC3*-HCM phenotypes.^29,33,52^

Similarly, heterozygous truncating variants in *DSP* are the major cause of ACM but phenotypes have not been consistently reported in heterozygous mouse models. Because complete *DSP* loss is fatal during fetal development, investigators have used conditional complete knock-out models.^17^ These models lead to an induction of total desmosomal loss that leads to rapid deterioration of heart function and does not replicate the human disorder. Strategies that mitigate complete failure in such a model may not necessarily translate to benefit in the heterozygous condition, limiting the fidelity of pre-clinical testing. In contrast, the haploinsufficient *Dsp* model developed here exhibits subepicardial cardiac fibrosis, a hallmark feature of ACM, without early decline in heart function, allowing future studies to decipher mechanisms and treatment strategies without the confounding effects of advanced cardiac remodeling.

Our quantitative modeling of mRNA versus protein abundance here revealed a steep, non-linear Hill relationship for both *Mybpc3* and *Dsp*. This sigmoidal dynamic reflects a saturable capacity for post-translational buffering.^24^ While mild transcript reductions (∼30-50%) are fully offset by decreased protein degradation rates, lowering transcript levels below critical thresholds (∼20-30%) overwhelms this buffering capacity, precipitating rapid protein loss and disease phenotypes. Our prior work using iPSC-CM models of *MYBPC3*-HCM implicated the Hsp70 chaperone system as a key mediator of post-translational compensation of cMyBP-C, and the Hsp70-Bag3 complex has also been identified as a direct regulator of cMyBP-C level.^12,53^ Availability of an in vivo model will allow future studies to further define these mechanisms. Additionally, Dsp protein level was excessively reduced at *Dsp* mRNA levels below ∼20% of normal. This pattern is broadly consistent with kinetic models of steady-state protein regulation in which translation efficiency itself scales non-linearly with mRNA abundance, such that low-abundance transcripts are translated less efficiently, allowing basal degradation to outpace protein synthesis.^54^

Generation of these haploinsufficient models will enable future studies to develop and test strategies to restore protein levels for both of these conditions. Such strategies could include either increasing production (e.g., gene replacement therapy or upregulation of transcription) or reducing degradation (e.g., inhibiting degradation pathways). CRISPRa has recently been advanced as a strategy to therapeutically upregulate gene expression with potential shown for multiple haploinsufficient conditions.^45,55–57^ In particular, CRISPRa and similar strategies circumvents the packaging limits for current gene replacement viral vectors. Specifically for *DSP*, we recently demonstrated the capability of transcriptional upregulation by CRISPRa to rescue haploinsufficiency and cell adhesion dysfunction in iPSC-CM cardiac tissue models.^34^ Targeting regulatory sequence elements is another promising approach with broad potential for haploinsufficient disorders.^58^ However, these and other approaches to restore gene expression stoichiometry will critically require in vivo testing for pre-clinical validation and optimization of correct gene dose levels. While new approach methodologies (NAMs), such as iPSC-derived cardiac tissue models, are an important initial platform for treatment development, we contend that in vivo models are essential for whole organ and system-level evaluations of these and related approaches. In addition to validation in two novel mouse models, we generated a compendium of SpliceAI predictions for haploinsufficiency genes using the MASC strategy to facilitate future model development for other haploinsufficiency mediated disorders. We found that SpliceAI predictions were highly correlated with in vivo allelic expression for our MASC designs. Our use of a canonical U2-type splice acceptor sequence upon which to base the MASC designs likely contributed to this predictable behavior. Although splicing can be highly dependent on local sequence context and tissue type^59^, SpliceAI predictions for MASC insertions replacing splice acceptor sites at constitutively spliced exons were consistent with broad generalizability across genes expressed in a range of tissue types. We did observe some variability that was dependent on the specific context sequence – e.g., some exons may exhibit a larger variance across the MASC variants, as was the case for *Ttn*. When implementing the technique for other genes, additional exons could be tested to identify optimal insertion sites that yield predictions with a range of splice modulation. Finally, this same approach could be useful to modulate gene expression in human cell models to create more robust disease phenotypes short of complete knock-out models. Consistent with this concept we have found that biallelic combinations of hypomorphic variants were similarly important for establishing robust disease models in both *MYBPC3*-HCM and *DSP*-ACM human iPSC-based cell models.^27,34^

In conclusion, we have developed a generalizable, titratable approach for modulating gene expression dose to more closely approximate haploinsufficiency levels observed in human disease. Using this approach, we generated two new mouse models that recapitulate key phenotypic features of human cardiomyopathy and identified potential utility across haploinsufficiency-associated genes more generally. Generation of these models will improve fidelity of mechanistic discovery and pre-clinical therapeutic development for these disorders. In particular, the MASC platform leaves native promoter/enhancer regulatory landscapes fully intact, making it uniquely suited for long-term physiological modeling and evaluating treatment strategies to restore normal gene expression levels.

## Methods

### Computational prediction of alternative splice cassette effects

All SpliceAI generated predictions were obtained by running SpliceAI version 1.3.1 via the python interface using a custom wrapper. Scores were calculated using mouse genomic context (mm39) with a scored distance of 500bp. Predictions for *Mybpc3* were generated for exon 4 (ENSMUST00000111430) with the human splice acceptor and exon 4 sequence (tgctcacagggtcaagctcagcagctctcaatggtcctacccctg) substituted for the corresponding mouse sequence (tcttttcagggtcagtctcggtaacccaggatggctcagctgcagagcatcagg). Variant effects as shown in Figure 1C were then tested in this humanized sequence context. Predictions for *Dsp* were generated for exon 14 (ENSMUST00000124830). Variant effects as shown in Figure 3H were then tested. Predictions for *Ttn* were generated for exons 81-88 on the N2-B isoform (ENSMUST00000011934). For predictions across haploinsufficiency genes, human genes defined as haploinsufficient were converted to mouse orthologs via Ensembl (release 115) and a transcript stable ID was obtained for each gene. The mm10 VastDB PSI TABLE was subset for genes of interest^46^ and VastDB event coordinates were converted to mm39 using the UCSC LiftOver tool.^60^ The GENCODE comprehensive gene annotation GTF file (GENCODE release M38) was used to define exons for each gene.^61^ To identify the exon with highest PSI across tissues, the following process was followed for each gene. First, VastDB information for the gene was filtered to remove uninformative splicing events (e.g. events with only a single coordinate, intron retention events). Samples with no splicing events that both met the minimum quality threshold and had PSI > 0 were also removed. For each exon in the transcript (excluding exon 1), if the exon coordinates were present in VastDB, an “inclusion score” was calculated defined as the number of samples with PSI > 90% divided by the total number of samples available for the gene. The exon with the highest inclusion score was chosen for downstream processing. If no exons passed the 90% threshold, then the threshold was iteratively lowered until an exon was found. The second exon was chosen for genes in which inclusion scores could not be calculated. Single exon genes were excluded. To test the effect of the MASC, the 33 bases 5’ of the splice acceptor site of the chosen exon for each gene was replaced with the sequence of each MASC variant and SpliceAI scores were obtained as described above.

### CRISPR-Cas9 editing in mice

CRISPR/Cas9 technology was used introduce mutations into *Mybpc3* (MGI:894710; ENSMUSG00000002100) and *Dsp* (MGI:109611, Ensembl: ENSMUSG00000054889). The CRISPOR algorithm was used to identify single guide RNA (sgRNA) targets with high predicted efficiencies (sgRNA protospacer sequences shown in **Supplemental Table 2**).^62,63^ Phosphorothioate modified sgRNAs (30 ng/ul per gRNA, Synthego) were complexed with wild type Cas9 protein (50 ng/ul, Synthego) to form ribonucleoprotein complexes (RNP).^64^ RNPs were microinjected into fertilized mouse eggs. Eggs were placed in culture until they developed into blastocysts. DNA was extracted from individual blastocysts for analysis. PCR with primers spanning the predicted cut site was used to generate amplicons for Sanger sequencing.^65^ Sequencing electropherograms of amplicons from individual blastocysts were evaluated to determine if small insertions/deletions caused by non-homologous endjoining (NHEJ) repair of chromosome breaks were present (primer sequences in **Supplemental Table 3**).^66^ The use of high specificity sgRNA and high fidelity Cas9 protein dramatically reduces the likelihood of off-target hits in mice.^67^ RNPs were mixed with a spot dialyzed synthetic short single stranded DNA donors (5 ng/ul, IDT) prior to microinjection into mouse zygotes for targeting MASC insertions at the splice acceptor sites for *Mybpc3* exon 4 and *Dsp* exon 14.^68^ The repair templates for each of the *Mybpc3* and *Dsp* MASC insertions were mixed in equimolar ratios to generate multiple knock-in alleles simultaneously in the same rounds of editing (**Supplemental Table 4**). The DNA donors were designed to replace the wild type sequence with the region containing the substitutions. Silent coding changes in the sgRNA binding sequence were included in the oligonucleotide to block cutting by Cas9 after repair of the chromosome by homology directed repair.^69^ RNPs were microinjected into mouse zygotes for *Dsp* exon 23 without a repair template to generate null alleles only to generate the *Dsp*^TV1k^^o/+^ model. The CRISPR reagents were microinjected into fertilized mouse eggs produced by mating superovulated C57BL/6:SJL (B6SJLF1) F1 Hybrids female mice (Jackson Laboratory stock no. 100012) with B6SJLF1 male mice as described.^70^ CRISPR/Cas9 microinjection of zygotes produced potential founder mice. PCR amplicons were Sanger sequenced (see **Supplemental Table 3**), and TOPO TA cloning was then performed on DNA from animals with evidence of the introduced changes to confirm the modified allele. G0 mice were backcrossed onto C57B6 mice for *Mybpc3* and onto DBA2J mice for *Dsp*. Phenotypes were assessed after >5 generations of backcrossing. The *Dsp^Ex^*^3d^*^el/+^* mouse model, generated as part of the NIH Knockout Mouse Project (KOMP), was obtained from the Mutant Mouse Resource and Research Center (MMRRC strain ID: *Dsp*^em1^(IMPC)^Mbp^/Mmucd).

### Echocardiography

Cardiac structure and function were assessed by transthoracic echocardiography. Mice were anesthetized according to the approved animal protocol using 3% isoflurane for induction, followed by 1–1.5% isoflurane for maintenance, titrated to maintain a heart rate >400 beats/min. Echocardiographic imaging was performed using a Vevo 2100 ultrasound system (FUJIFILM VisualSonics) equipped with an MS550D 40-MHz transducer. Two-dimensional B-mode images were acquired in parasternal long-axis, parasternal short-axis, and apical views. Left ventricular (LV) systolic function was assessed by calculating ejection fraction from the parasternal long-axis view as: EF (%) = [(LVEDD³ − LVESD³)/LVEDD³] × 100, where LVEDD and LVESD represent LV internal diameter at end-diastole and end-systole, respectively. LV mass was calculated as: LV mass = 1.053 × [(IVSd + LVEDD + LVPWd)³ − LVEDD³]/1.25, where IVSd and LVPWd represent interventricular septal and LV posterior wall thickness at end-diastole, respectively. Pulsed-wave Doppler recordings of mitral inflow were obtained from the apical view and used to measure isovolumic relaxation time (IVRT) as an index of diastolic function and isovolumic contraction time (IVCT) and aortic ejection time (ET) as indices of systolic function. Measurements were obtained from representative cardiac cycles under stable physiologic conditions.

### Mouse cardiac tissue collection and processing

Mice were euthanized by carbon dioxide inhalation according to the approved animal protocol. Immediately following euthanasia, hearts were perfused with ice-cold phosphate-buffered saline (PBS) through the ventricular apex to remove residual blood. Hearts were then excised, rinsed in ice-cold PBS, and dissected for downstream histologic, proteomic, and RNA sequencing analyses. Tissue designated for histologic analysis was either fixed for subsequent paraffin embedding in 10% neutral buffered formalin for 24 hours or embedded in OCT (Tissue Tek) for frozen sectioning. Tissue designated for proteomic analysis was snap-frozen on dry ice and stored at −80°C until processing. Tissue designated for RNA sequencing was immersed in RNAlater stabilization solution and incubated at 4°C overnight. The following day, tissue was removed from RNAlater and stored at −80°C until RNA isolation and downstream processing.

### RNA sequencing and analysis

Cardiac tissues were homogenized on ice in RLT buffer supplemented with β-mercaptoethanol, and total RNA was isolated using the RNeasy Kit (Qiagen). RNA concentration and purity were assessed by NanoDrop spectrophotometry, and RNA integrity was evaluated before library preparation. Samples had RNA concentrations >25 ng/µL, A260/A280 ratios >1.9, and RNA integrity numbers >8.0. Stranded, barcoded libraries were prepared and multiplexed by the University of Michigan DNA Sequencing Core, followed by 151-bp paired-end sequencing on an Illumina NovaSeq X Plus. Demultiplexed FASTQ files were generated using BCL Convert. A Snakemake-managed workflow was used to trim reads with Cutadapt, assess read quality with FastQC, align reads to the mouse GRCm38 reference genome (Ensembl release 102) using STAR with ENCODE-recommended RNA-seq parameters, and estimate gene-level counts using RSEM. Pipeline quality-control metrics were aggregated using MultiQC.

Lowly expressed genes were excluded before differential-expression analysis. Raw gene-level counts were normalized by size-factor estimation and analyzed in DESeq2 using gene-specific dispersion estimates, negative binomial generalized linear models, and Wald tests. Differential-expression results were visualized with EnhancedVolcano. Gene Ontology over-representation analysis was performed with clusterProfiler using genes with a DESeq2-adjusted P < 0.05 and absolute log2 fold change >0.2, default hypergeometric test and Benjamini–Hochberg correction were applied. clusterProfiler’s gseGO and gseKEGG functions were utilized for gene set enrichment analysis. Enrichment results were visualized using enrichplot and ggplot2. Transcriptomic data were additionally analyzed and visualized using Advaita Bioinformatics’ iPathwayGuide. Sashimi plots were generated using the Integrative Genomics Viewer. RNA-seq data will be deposited in the NCBI Gene Expression Omnibus (accession pending).

### Mybpc3 splice-junction analysis

To evaluate alternative splicing of *Mybpc3* variants, a single-contig *Mybpc3* reference sequence containing the humanized exon 4 sequence was generated from the GRCm38 chr2 locus. The endogenous sequence was replaced with the 72-nucleotide humanized sequence, which was 9 nucleotides shorter than the corresponding mouse sequence and included a 9-nucleotide intronic prefix. A STAR index was constructed and reads mapping to the *Mybpc3* locus were extracted from the whole-genome BAM files with SAMtools. The reference sequence, annotation, and aligned BAM files were loaded into the Integrative Genomics Viewer, and splice-junction usage was visualized using Sashimi plots. Reads spanning junctions the canonical and alternative splice acceptor sites were quantified. Alternative splice-junction usage was calculated using a correction factor to account for 85% steady state NMD of alternatively spliced transcripts (determined from absolute abundance levels of truncating variant transcripts in homozygous null *Mybpc3* mice) with the following equation: (alternative-junction reads/0.15)/(alternative-junction reads/0.15 + canonical-junction reads) × 100.

### Liquid chromatography mass spectrometry (LCMS)

A 1-2 mg piece of the left ventricle of each heart was solubilized with Rapigest (Waters), the proteins were reduced with dithiothreitol, alkylated with iodoacetamide (Acros Organics), and digested to peptides using trypsin (Promega) as described.^14^ The Rapigest was cleaved, and peptides extracted in 0.1% trifluoracetic acid as described.^14^ Peptides were separated by injection onto an XSelect HSS T3 column (3.5 μm, 1.0 mm × 150 mm) (Waters Corporation) attached to an UltiMate 3000 ultra-high pressure liquid chromatography (UHPLC) system (Dionex). The UHPLC effluent was directly infused into a Q Exactive Hybrid Quadrupole-Orbitrap mass spectrometer by electrospray ionization (Thermo Fisher Scientific). Data were collected in data dependent MS/MS mode with the top five most abundant ions being selected for fragmentation, as previously described.^14^ The .raw LCMS files were searched using Sequest HT against the Mus musculus UniProt reference proteome (74,085 sequences; downloaded 02/09/2015) in Thermo Proteome Discoverer (v2.2.0.388) as described.^14^ Protein abundances were determined from the average abundance of the LC peak areas of the top 5 ionizing peptides per protein. Either the average ratio or the ratio normalized to the most abundant protein in each group were reported.

### Histological Analyses

Cardiac tissues were fixed in formalin for 24 hours at 4°C and subsequently transferred to 70% ethanol for storage. Samples were then processed and embedded in paraffin wax using a Leica Histocore Arcadia embedding center. Paraffin-embedded blocks were sectioned at a thickness of 7 µm using a Leica RM2255 semi-automated microtome and mounted to slides. To evaluate myocardial fibrosis, sections were stained with picrosirius red and fast green. Sections were first incubated in 0.04% fast green for 15 minutes. Following a wash with distilled water, the slides were incubated in a solution of 0.1% fast green and 0.04% sirius red in saturated picric acid for 30 minutes to differentially stain the collagenous tissue and healthy myocardium. Stained slides were digitally scanned using a Leica Aperio AT2® whole slide scanner at 20x magnification. Quantitative analysis of the images was performed using ImageJ software to assess and calculate the ratio of fibrosis (indicated by picrosirius red staining) to the total myocardium (indicated by fast green staining).

### Cryosectioning and Immunofluorescence Staining

Harvested hearts were immediately embedded in optimal cutting temperature (OCT) compound and flash-frozen. Frozen tissue blocks were sectioned at a thickness of 7 µm using a cryostat (Leica, CM3050 S) and mounted to slides. For immunofluorescence staining, frozen sections were fixed in cold 4% paraformaldehyde (PFA) for 5 minutes, followed by permeabilization with 0.1% Triton X-100 for 5 minutes. To minimize non-specific antibody binding, the tissue sections were blocked in 5% normal goat serum for 45 minutes at room temperature. Sections were then incubated overnight at 4°C with primary antibodies diluted in 5% goat serum. The following primary antibodies and markers were utilized depending on the experimental panel: mouse anti-alpha-actinin (1:1000, Sigma A7811), anti-CD45 monoclonal antibody (1:100, Invitrogen, Cat# 50-125-87), rabbit anti-Col1A1 polyclonal antibody (1:100, Invitrogen, Cat# PA5-29569), anti-CCR2 (1:100, Biolegend, Cat#105619), wheat germ agglutinin (WGA) – Aat Bioquest, Cat# 25512. Following primary incubation, sections were washed and incubated with the corresponding fluorophore-conjugated secondary antibodies (1:1000) for 45 minutes at room temperature. Secondary antibodies included goat anti-mouse Alexa Flour 647 (Invitrogen, A21235), goat anti-rabbit Alexa Fluor 568 (Invitrogen, Cat# A11036), goat anti-rat Alexa Fluor 488 (Invitrogen, A48262). Nuclei were counterstained using ProLong™ Diamond Antifade Mountant with DAPI (Invitrogen, Cat# P36962) during slide mounting. Fluorescence images were obtained using a Nikon Eclipse Ti-E inverted microscope at 40x magnification.

## Acknowledgements

We acknowledge Judy Miller and Janet Miller-Monfils for their generous support. We acknowledge Thomas L. Saunders, Zachary T. Freeman, Elizabeth Hughes, Hongali Zhang, Wanda Filipiak, Galina Gavrilina, and the Transgenic Animal Model Core (RRID:SCR_000776) of the University of Michigan’s Biomedical Research Core Facilities for their assistance in design and production of transgenic mice. Research reported in this publication was supported by the University of Michigan Transgenic Animal Model Core and the Biomedical Research Core Facilities.

