## Supplemental Figures for "Genomic Engineering of Gene Dosage: A Generalizable Framework for Modeling Haploinsufficiency-Mediated Human Disorders through Splicing Modulation"

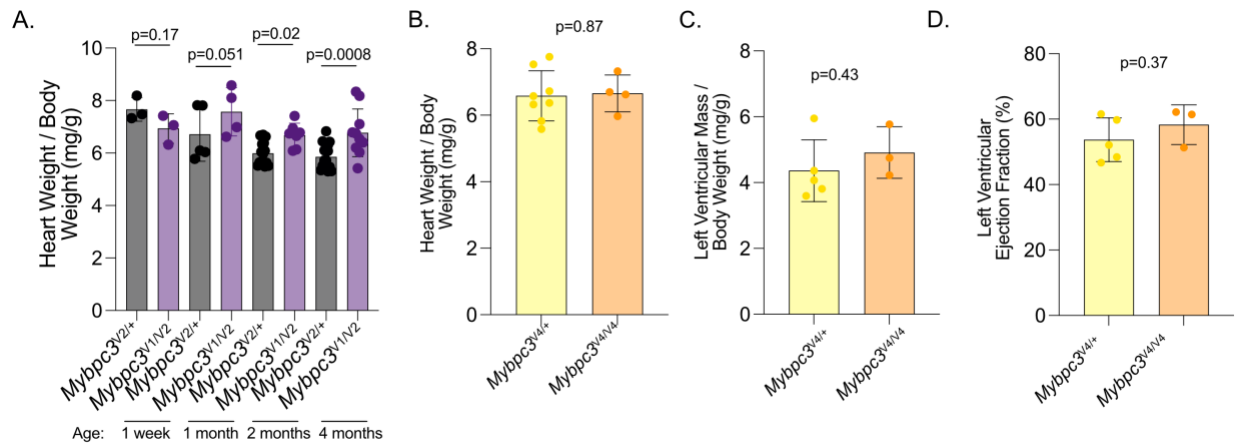

**Supplemental Figure 1. MyBPC-C haploinsufficiency causes increased cardiac mass, while mice with a humanized exon 4 but without haploinsufficiency do not exhibit evidence of HCM. A.** *Mybpc3<sup>V1/V2</sup>* mice exhibit an increase in heart mass to body weight by 2 months of age compared to littermate controls without c-MyBP-C haploinsufficiency (*Mybpc3<sup>V2/+</sup>* mice). **B.** *Mybpc3<sup>V4/V4</sup>* mice, which have the same humanized exon 4 and missense variant as in *Mybpc3<sup>V1/V2</sup>* mice but normal splicing and normal cMyBP-C protein level, do not exhibit an increase in heart weight to body weight ratio at 2 months of age (*Mybpc3<sup>V4/V4</sup>* n=4 versus *Mybpc3<sup>V4/+</sup>* n=8 littermate controls). **C-D.** Echocardiograms at 1 month of age demonstrate no hypertrophy (C) or change in LV ejection fraction (D) in *Mybpc3<sup>V4/V4</sup>* mice (*Mybpc3<sup>V4/V4</sup>* n=3 versus *Mybpc3<sup>V4/+</sup>* n=5 littermate controls).

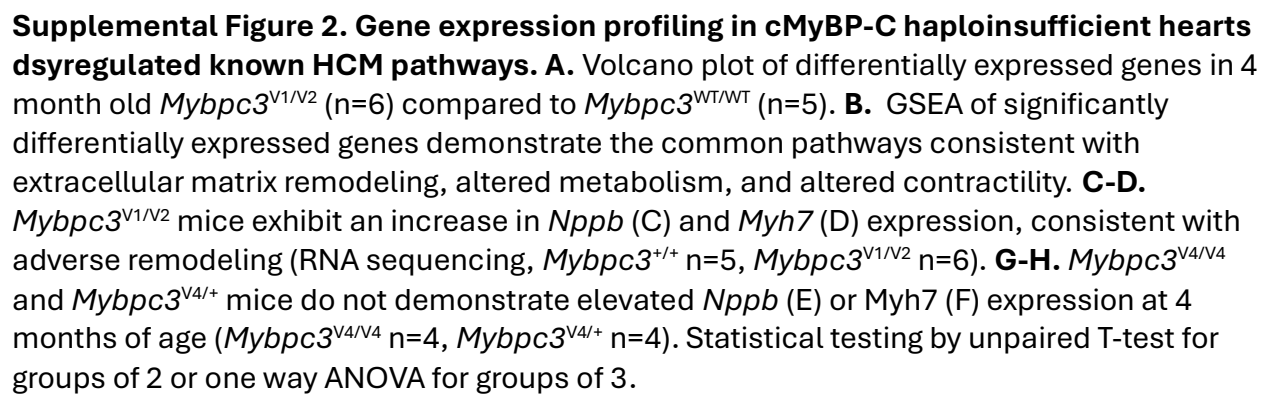
